# Multiplexed single-molecule analysis of human telomerase synthesizing DNA

**DOI:** 10.1101/2021.06.25.449954

**Authors:** Jendrik Hentschel, Junhong Choi, Clive R. Bagshaw, Christopher P. Lapointe, Jinfan Wang, Linnea I. Jansson, Mareike Badstuebner, Joseph D. Puglisi, Michael D. Stone

## Abstract

Genomic stability in proliferating cells critically depends on telomere maintenance by telomerase reverse transcriptase. Here we developed a real-time single-molecule RNA sequencing approach that visualizes telomerase catalysis and structural dynamics at single-nucleotide resolution using FRET and zero-mode waveguides. The method permits direct detection of dynamic steps and structural states throughout the telomerase catalytic cycle and can be generalized to other nucleic acid polymerase systems.

## Main

Telomere homeostasis is critical to tissue development and regeneration. The telomerase enzyme synthesizes the repetitive telomere DNA that, together with telomere-binding proteins, protects linear chromosome ends.^1,2^ Enzyme dysfunction causes premature aging diseases, while aberrantly upregulated telomerase activity confers replicative immortality to most cancers.^3,4^ Improved understanding of telomerase structure and function would have direct biomedical significance.

Telomerase is a ribonucleoprotein (RNP) reverse transcriptase (RT) that uses an integral RNA-template to synthesize telomeric DNA-repeats (Fig. 1). In humans, the telomerase RNA (TR) template sequence 3’-rC1-rC2-rA3-rA4-rU5-rC6-5’ specifies the hexameric DNA-repeat sequence GGTTAG.^1^ The telomerase enzyme (E) exhibits two types of processivity. Nucleotide-addition processivity (NAP) describes the incorporation of up to six nucleotides (N) into the DNA substrate (D) releasing pyrophosphate (P) upon nucleotide hydrolysis. NAP is associated with single-nucleotide translocation steps of the TR-template in the active site.

**Fig. 1:**
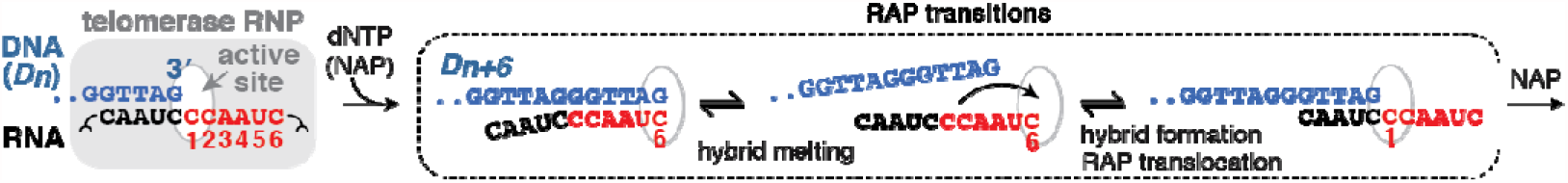
The human telomerase catalytic cycle. Schematic of the telomerase RNP reverse transcription reaction. Nucleotide addition occurs in hexameric units followed by DNA/RNA translocation. RNA template positions are numbered in red.

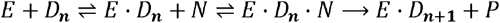

Repeat-addition processivity (RAP) describes the addition of multiple repeats to a single DNA-substrate. RAP consequently requires larger-scale translocation steps to reposition the TR-template in the active site from position rC6 to rC1 to enable another round of nucleotide addition (Fig. 1).^5^ The structural rearrangements required to promote RAP may be described by a dynamic equilibrium:

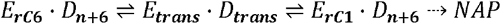

*E*_*trans*_ · *D*_*trans*_ represents a transient intermediate state characterized by a stable tether between telomerase ‘anchor-sites’ as well as a dynamic DNA-product during melting of the DNA/RNA hybrid and subsequent re-priming for the next round of telomere repeat synthesis.^6,7^

Established methods to investigate telomerase activity are primer-extension assays that monitor the growing DNA chain.^8^ Using sequencing gels and radiography, ensemble telomerase reactions can be analyzed with nucleotide resolution (*D*_*n*_ vs *Dn*_+ 1_) and enable global kinetic modeling of telomerase RAP (Fig. 2a).^9^ The RAP translocation process exhibits rate-limiting step(s), at which extension products accumulate. Repeat synthesis by telomerase is slow, with an average time span of several minutes per individual repeat.^9^ Telomerase exhibits relatively high Km values for nucleotide binding, and requires micro-molar dNTP concentrations for robust activity.^10^ More recently, single-molecule approaches probed the growing DNA product in real time (*E* ·*D*_*n*_ vs *E* · *D*_*n* + *x*_), but mechanistic and kinetic insights into NAP and RAP cycles were obstructed by DNA structural dynamics.^9,11,12^ Extensive cryo-EM efforts have provided insight into the architecture of the ciliate and human telomerase RNP, including higher-resolution snapshots of the DNA-RNA hybrid in the active site.^13,14^ These biochemical and structural studies highlight the large degree of conformational heterogeneity intrinsic to telomerase; however, understanding how these structural dynamics relate to function requires direct visualization of actively-extending telomerase enzymes.

**Fig. 2:**
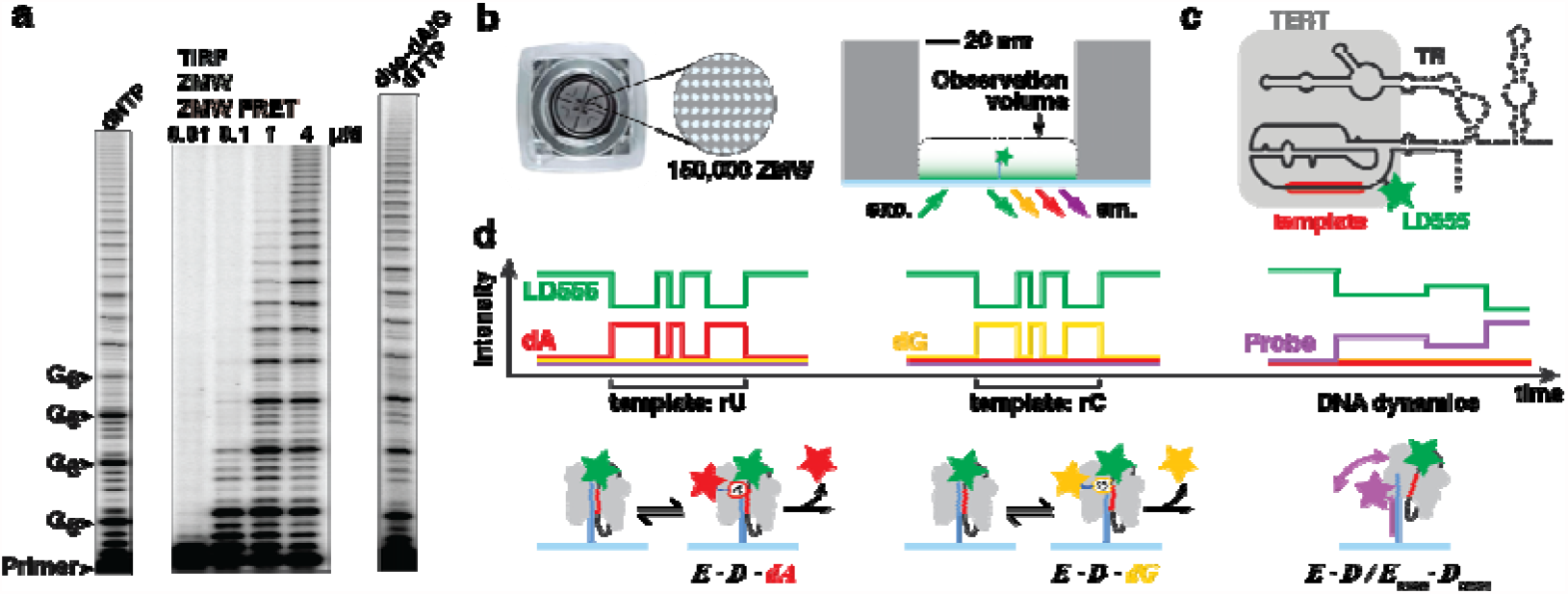
The human telomerase ZMW-FRET approach. **a**. Telomerase primer extension assays resolved on sequencing gels. Telomerase requires micro-molar nucleotide concentrations. An approximated dynamic range of single-molecule methods is given above the gel. Telomerase utilizes dye-dN6P. **b**. Geometry of zero-mode waveguide array chips used for telomerase multicolor ZMW-FRET. **c**. Schematic of human telomerase reverse transcriptase (TERT) and the telomerase RNA (TR, solid and dashed lines). LD555 at the U42 dye-labeling site is highlighted. The catalytic core is formed by TERT and essential domains of TR (solid lines). **d**. Principal signals expected from multiplexed ZMW-FRET. Dye-coupled hexaphosphate nucleotide analogs dA6P (red) and dG6P (yellow) interact reversibly with telomerase. Eventual irreversible incorporation releases the dye moiety. At right, a Cy5.5-labeled DNA-probe (magenta) can bind to nascent telomere DNA and reports on DNA structural dynamics. Cartoon schematic (lower panel) depicting telomerase enzyme (E) and substrate DNA (D) complexes.

To address these limitations, we developed a single-molecule real-time RNA sequencing approach that visualizes telomerase-nucleotide complexes directly (*E* · *D*_*n*_· + *N*) and permits concurrent detection of functional structural dynamics during the telomere synthesis reaction. The method overcomes several technical challenges in single-molecule studies, facilitating experiments that (i) increase the concentration of dye-coupled nucleotides to physiologically relevant concentrations, (ii) differentiate enzyme-bound nucleotides from unspecific fluorescence signals, and (iii) enable real-time identification of conformational dynamics that govern enzyme function (Supplementary Fig. 1). The technique builds on the Pacific Biosciences SMRT-sequencing approach (SMRT: single-molecule real-time, Fig. 2b and Supplementary Fig. 1a).^15^ However, a key differentiating principle of our approach is the use of Förster Resonance Energy Transfer (FRET) to measure the distance-dependent transfer of energy between dye-conjugated nucleotide analogs and an enzyme-linked fluorescent dye. In contrast to the direct excitation of fluorescent components across the visible light spectrum employed in SMRT sequencing, the use of FRET permits experiments to be conducted at higher, near physiological, concentrations of dye-conjugated nucleotide analogs (Fig. 2b-d and Supplementary Fig. 1)^16^. Dye-coupled dATP and dGTP nucleotide analogs serve as FRET acceptor molecules, and report on rU and rC template positions upon binding to telomerase (Fig. 1, 2d and Supplementary Fig. 2b). We reason that the combination of unmodified dTTP with color-coded dA/dG analogs produces unambiguous telomeric FRET patterns, while delineating the start and end-point of RAP transitions in real time (<u>GG</u>TT<u>AG</u> – RAP – <u>GG</u>TT<u>AG</u>). The phosphate-linked dye-moieties are released upon nucleotide incorporation, yielding a natural DNA product that does not interfere with downstream functions (Supplementary Fig. 2b).^15^ In a multiplexed FRET approach, DNA structural dynamics can then be visualized in parallel using a spectrally distinct dye-labeled DNA-probe bound to the nascent telomere DNA (Fig. 2d).

To visualize (*E* · *Dn*· *N*) complexes in real time we coupled a ‘self-healing’ FRET-donor fluorescence dye (LD555) to the TR subunit at residue U42 (Fig. 2c).^17,18^ Structural models suggest that this established labeling site is within FRET range of the nucleotide binding pocket (Supplementary Fig. 3a).^14,19^ To facilitate site-specific dye-labeling, the catalytic TERT-protein (‘Telomerase Reverse Transcriptase’) was reconstituted with essential domains of telomerase RNA *in-vitro* (Fig. 2c). This core-complex of the telomerase RNP is structurally characterized and shows comparable activity to the telomerase holoenzyme isolated from human cells (Supplementary Fig. 3b).^9,14,19^ Dye-coupled hexaphosphate-aminohexyl dA and dG nucleotide analogs (dye-dN6P) were synthesized at PacBio and coupled to fluorescence dyes following established protocols (Supplementary Fig. 2b).^20^ Telomerase incorporates both types of analogs in ensemble primer-extension assays with comparable activity when compared to unlabeled dNTP (Fig. 2a).

To observe telomerase activity directly in real time, LD555-telomerase was bound to a biotinylated (TTAGGG)_3_ DNA-primer and immobilized inside individual zero-mode waveguide (ZMW) wells on PacBio array-chips (Fig. 2b). Previous work has demonstrated the utility of a ZMW-compatible PacBio RSII real-time sequencer (RS) customized for single-molecule fluorescence spectroscopy (Supplementary Fig. 2a).^21^ Chips were supplied with imaging mix containing dye-d(A/G)6P and mounted on the RS. Four-color emission traces from each ZMW were recorded for 1800 seconds at 10 Hz detector frame-rate (Fig. 3a). To initiate telomerase primer extension in the presence of dye-d (A/G)6P, dTTP was added to the chip by automated pipetting during data acquisition. The low background signal afforded by a combination of ZMW-confinement with single laser-line excitation at 532 nm allowed for dye-dN6P concentrations that support robust telomerase activity (3.5 µM dG6P, 8 µM dA6P, 10 µM dTTP, Fig. 2a).

**Fig. 3:**
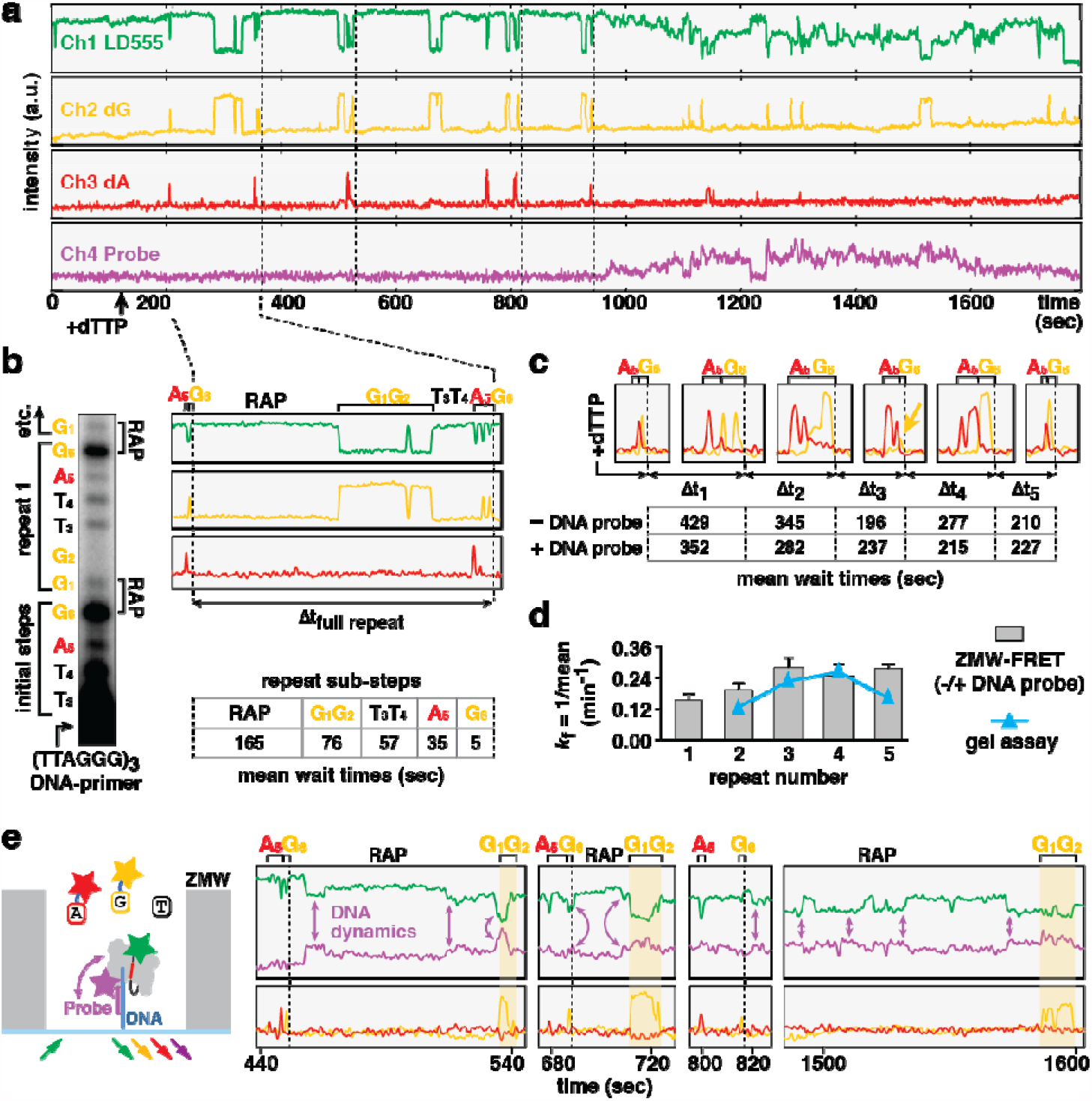
Real-time ZMW-FRET analysis of human telomerase. **a**. Four-color trace from a single telomerase molecule. Detection channels are shown separately for clarity. Dashed lines separate discernible repeat patterns (b),(c). **b**. At left, cropped gel lane visualizes early primer extension and informs expectations for a valid ZMW-FRET assay. Sequence and numbers indicate telomeric DNA-repeats and the respective template position for each nucleotide. At right, example of a repetitive FRET pattern consistent with a telomeric DNA repeat. Repeat sub-steps corresponding to telomerase RAP and NAP are indicated. The mean duration of each respective sub-step is given below, n_RAP_=219, n_G1G2_=373, n_T3T4_=153, n_A5_=577, n_G6_=576. **c**. A5/G6 motifs enlarged from trace in (a). G6 incorporations as repeat-end signatures are indicated (dashed lines) and yield repeat wait times for kinetic analysis (Δt), cf. Supplementary Fig. 8. Below, mean wait times for individual repeats derived from two RS datasets. **d**. Single-step forward rate constants are determined from mean wait times for individual repeats (c). The values are consistent with telomerase ensemble assays. **e**. Example trace of multiplexed ZMW-FRET visualizing DNA-dynamics (arrows highlight selected events). Time axis is broken to enlarge regions of interest, cf. Supplementary Fig. 5. Ch1/4 and Ch2/3 are superposed.

To probe telomerase-substrate conformational changes during active telomerase incorporation in real time, we performed separate experiments in the presence of Cy5.5-labeled DNA-oligonucleotides that are complementary to 2.5 telomeric repeats (Fig. 2d and Fig. 3a). ^9,10^ We have previously shown that individual DNA-probes anneal to nascent telomeric DNA within FRET-range of a U42-coupled donor dye.^9^

We obtained two datasets of 876 and 2009 traces (-/+ DNA-probe, respectively) that contained discernible FRET-events between LD555-telomerase and dye-dG6P (*E* · *Dn* · *dG*, yellow) or dye-dA6P (*E* · *Dn* · *dA*, red) (Fig. 3a and Supplementary Fig. 4-6). ZMW fluorescence emission exhibited spectral bleed-through into neighboring detection channels.^21^ Bleed-through and background-fluorescence were corrected by signal subtraction (Supplementary Fig. 7a). The source channels for specific FRET events could unambiguously be assigned. Non-cognate nucleotide sampling events on the microsecond timescale were not resolved on the timescale of the measurement (100 ms per frame). However, signal clusters from association and dissociation of more stably-bound cognate nucleotides could be visualized (*E* · *Dn* + *N* ⇌ *E* · *Dn* · *N*) (Fig. 2e and Supplementary Fig. 7b). These clusters report on the identity of a given template position, as has been described in ZMW-analysis of HIV reverse transcriptase.^22^

The ZMW-FRET approach presented here provides means to unravel complex biological problems in real time through multiplexed and quantitative single-molecule analysis. Certain features of these proof-of-concept experiments were expected and serve to validate the approach. Nucleotide addition to the telomerase primer (TTAGGG)3 was expected to yield FRET-signals in the red and yellow channels as the initial repeat was being completed (template positions rA3, rA4, rU5 and rC6) (Fig. 3b). This initial pattern was observed in ∼90% of all traces. Subsequent FRET-signals were expected to adhere to full telomeric repeat cycles, separated by RAP-translocation pauses. In a total of 817 representative traces across both datasets, we consistently identified 1134 repeat patterns extending beyond the initial A5/G6 signals. Analysis of each repeated pattern confirmed the presence of distinct sub-steps: (i) a ‘dark’ RAP-translocation phase, (ii) G1/G2-FRET-clusters, (iii) a ‘dark’ T3/T4-phase, and (iv) a A5/G6-FRET-cluster concluding the repeat (Fig. 3b and Supplementary Fig. 8a).

To validate the real-time measurement of processive telomere repeat synthesis, we extracted repeat wait times between individual G6-incorporation events and calculated single-step forward rate constants (Fig. 3c and Supplementary Fig. 8b). The obtained values from both independent datasets were consistent with corresponding rate constants derived from kinetic modeling of telomerase ensemble assays, further supporting the robust telomerase activity in ZMW-FRET (Fig. 3d).^9^

Consistent with prior biochemical analysis of ensemble telomerase activity, the RAP-translocation pause was the slowest sub-step within an individual repeat cycle with a mean duration of ∼165 seconds (Fig. 3b). An unexpected feature of our data was that G6-incorporation occurred relatively fast (∼5 sec), whereas the G1/G2-FRET-clusters exhibited a substantially increased mean duration (∼76 sec, Fig. 3b and Supplementary Fig. 6, 8a). The prolonged G1/G2 clusters are unlikely to be explained by the simple combination of two successive G-incorporation events, and therefore suggest the existence of a long-lived state of the enzyme that results in slow dG incorporation kinetics at the beginning of each telomeric DNA repeat (Supplementary Fig. 9). Although the precise nature of this sub-step remains to be elucidated, it is plausible that additional RAP-related events occur after template repositioning and G1-binding, including re-formation of the RNA/DNA hybrid, re-priming of the DNA 3’-end, and active-site closure (Supplementary Fig. 9).^7^

Our approach reveals telomerase-template dynamics during telomere synthesis. In the presence of Cy5.5-DNA-probe, 605 of 2009 traces displayed FRET between LD555-telomerase and the Cy5.5-labeled probe (magenta) bound to the nascent telomere DNA product. This observation represented a direct measurement of product DNA-dynamics resolved concurrently with nucleotide-addition cycles (Fig. 3e and Supplementary Fig. 5).

Interestingly, we identified repeated instances of a Cy5.5-FRET increase preceding the onset of G1/G2-clusters, suggesting a mechanism by which DNA-dynamics must precede re-positioning of the TR-template to allow for G1-binding (Fig. 3d). This result is preliminary and will require further investigation to substantiate. However, the ability to resolve product DNA-dynamics at specific stages of telomerase repeat synthesis provides a powerful illustration of the information-rich data that can be obtained by combining SMRT-sequencing with single-molecule FRET in the study of nucleic acid polymerases.

**Acknowledgements**

We are grateful to Lubomir Sebo and Jonas Korlach of Pacific Biosciences for synthesizing and providing dye-coupled nucleotide analogs for this study. We thank Carol Greider for discussion of the results. J.H. was supported by a Swiss National Science Foundation (SNSF) Early Postdoc.Mobility ellowship (P2EZP3_181605). J.C. was supported by a Stanford Bio-x Fellowship and is a Howard Hughes Medical Institute Fellow of the Damon Runyon Cancer Research Foundation (DRG-#2403-20). C.P.L. is a Damon Runyon Fellow supported by the Damon Runyon Cancer Research Foundation (DRG-#2321-18), and J.W. was supported by a postdoctoral scholarship from the Knut and Alice Wallenberg Foundation (KAW 2015.0406). This work was supported by National Institutes of Health Grants F99CA212439 (to L.I.J.) and R01GM095850 (to M.D.S.), as well as grants GM51266, GM011378 and AI150464 to J.D.P

## Methods

See Online Methods section below.

**Supplementary Fig. 1.**
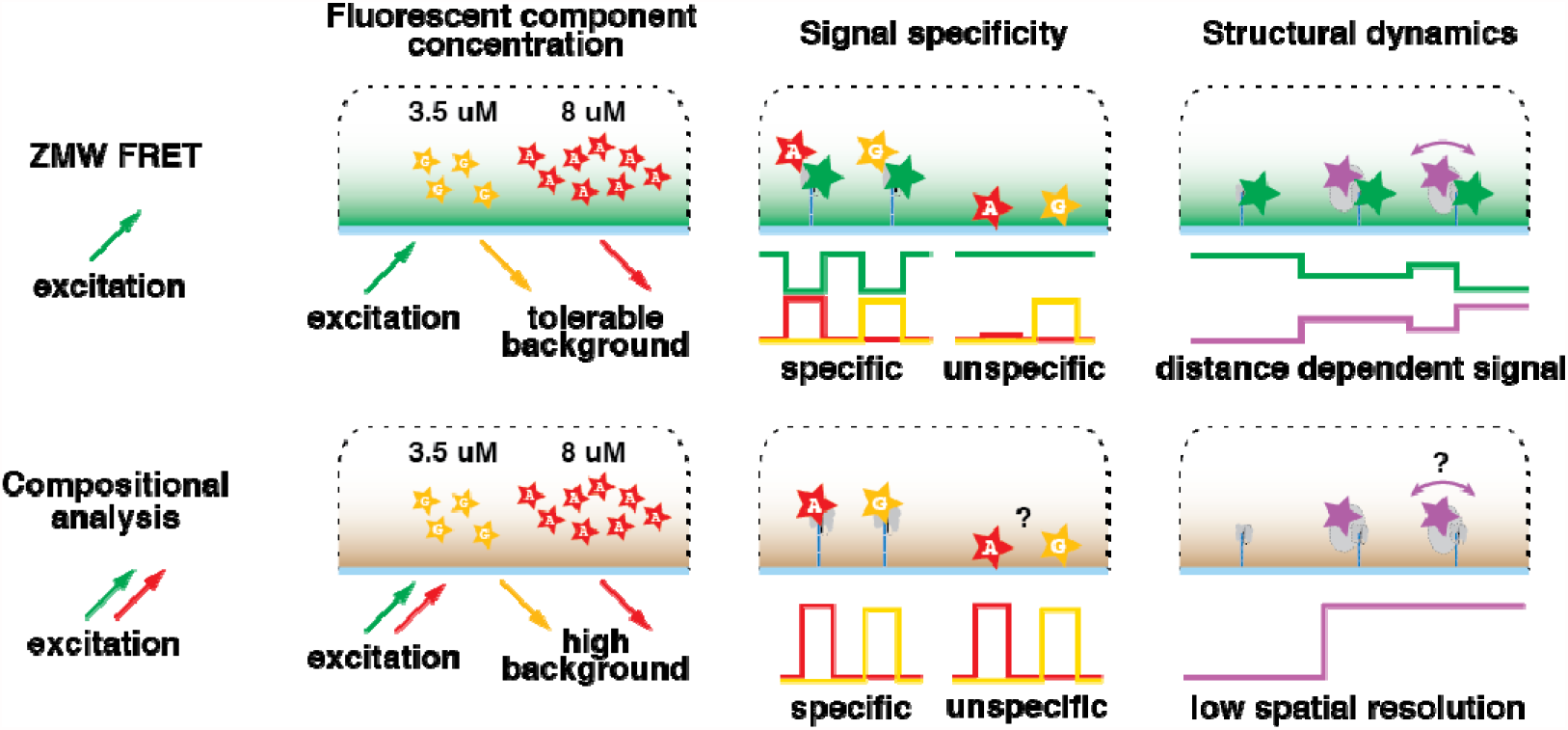
Specific advantages of ZMW-FRET over SMRT-sequencing when applied to mechanistic studies of telomerase reverse transcription or other biological systems.

**Supplementary Fig. 2.**
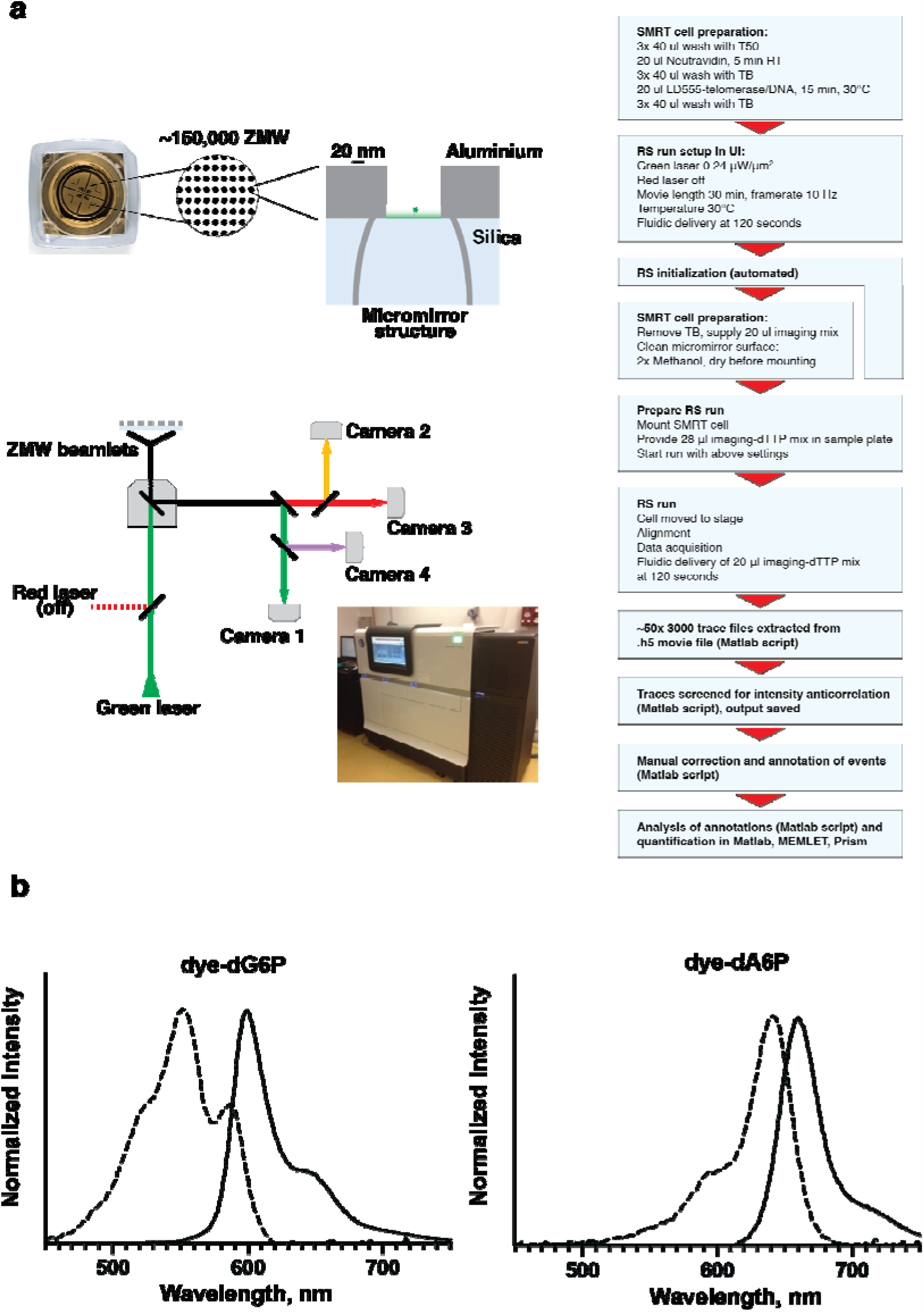
a. Principle setup of ZMW chips and the PacBio RS II instrument used for telomerase ZMW-FRET real-time analysis. **b**. Excitation (dashed lines) and emission spectra of dye-dG6P and dye-dA6P. Absorption data collection on Nanodrop UV spectrophotometer and emission data collection on a Varian fluorimeter.

**Supplementary Fig. 3.**
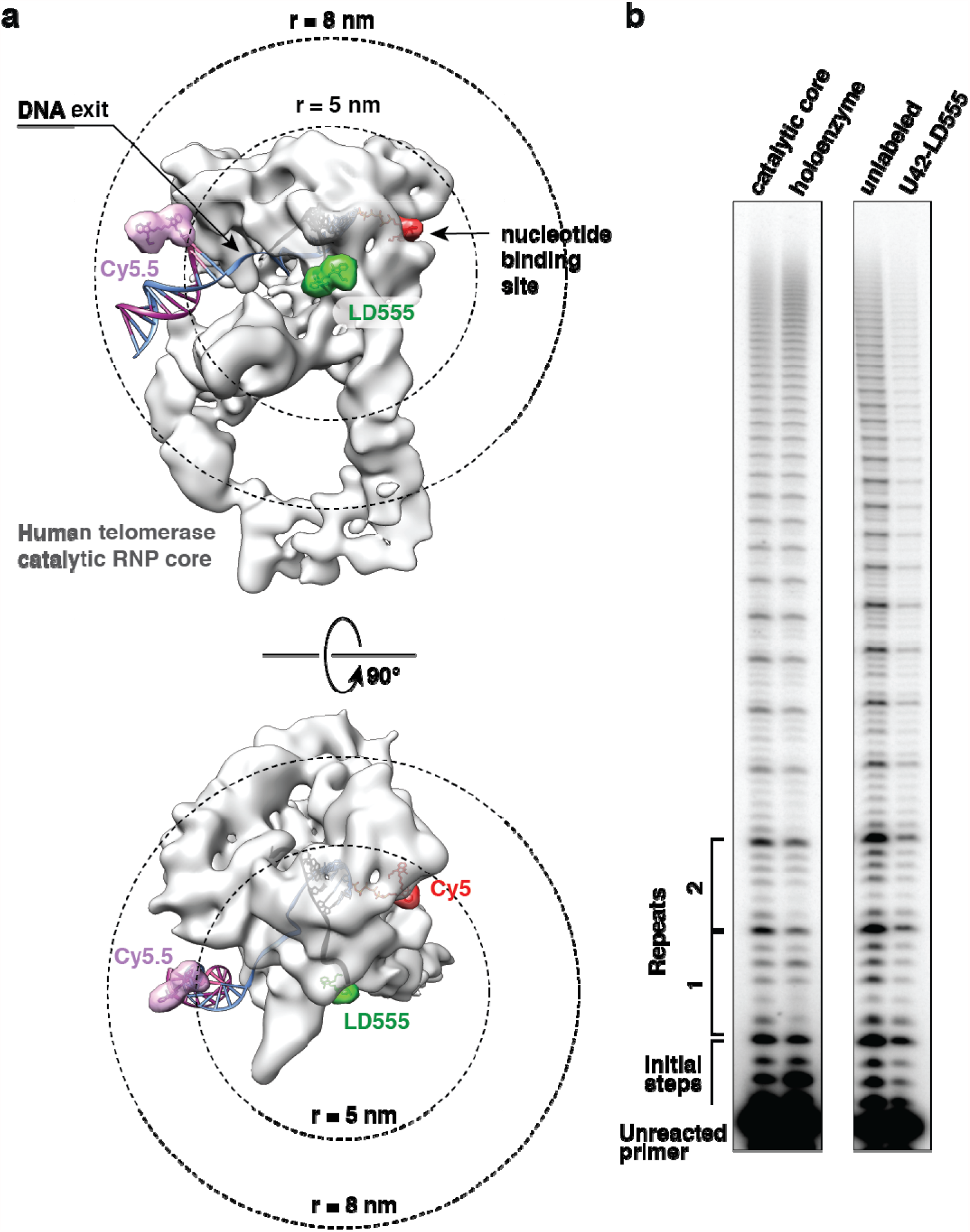
Human telomerase catalytic core. **a**. The cryo EM structure (EMDB 7518) approximates the minimal telomerase complex used in this study. Telomeric DNA (33 nt) is in blue, telomerase RNA template region in black. The nucleotide binding pocket is occupied with a coordinate model of dA6P-aminohexyl coupled to Cy5 (red). The U42 labeling site and approximate position of LD555 are indicated with a simulated density map of a standard Cy3 fluorescence dye moiety (green). A Cy5.5-DNA probe is annealed to nascent telomere DNA in a possible orientation outside the DNA exit cavity of telomerase. FRET radii around LD555 are indicated by dashed circles. Bottom, top view. **b**. Left, primer extension assays of the *in-vitro* reconstituted telomerase catalytic core compared to telomerase holoenzyme isolated from HEK293T cells. Right, primer extension assay of unlabeled telomerase catalytic core and LD555-telomerase catalytic core. Note that dye-labeled enzyme is reconstituted with limiting amounts of LD555-hTR fragments leading to lower enzyme yields and product band intensities.

**Supplementary Fig. 4.**
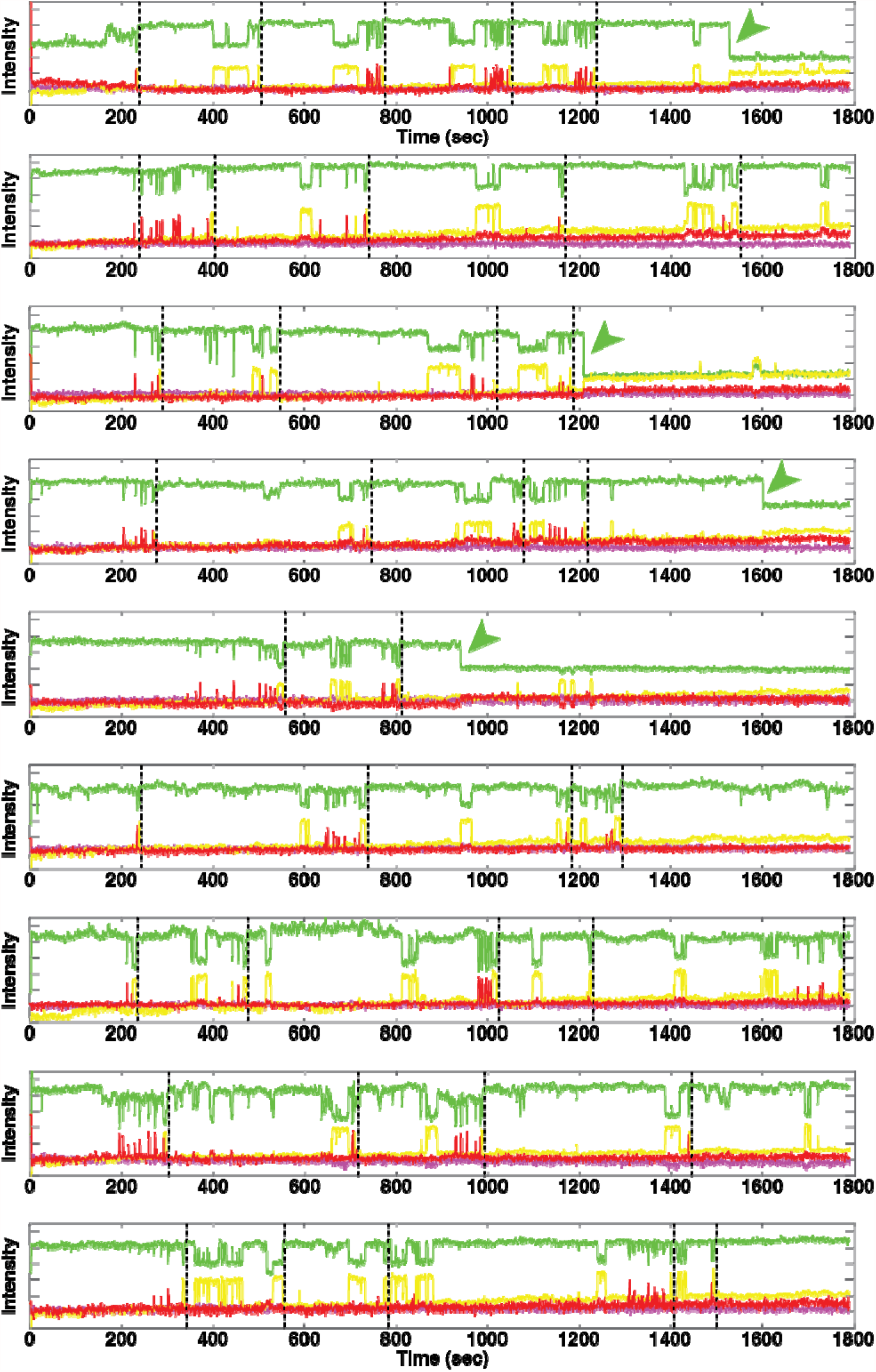
Example traces visualizing real-time telomerase activity by ZMW-FRET in absence of DNA-probe oligonucleotide. Dashed lines indicate G6 incorporation events preceding a RAP phase, cf. Fig. 3a-c. Loss of LD555 donor fluorescence (green arrows) represents DNA dissociation from telomerase or dye photo-bleaching. An offset is applied to the baseline of LD555 for clarity.

**Supplementary Fig. 5.**
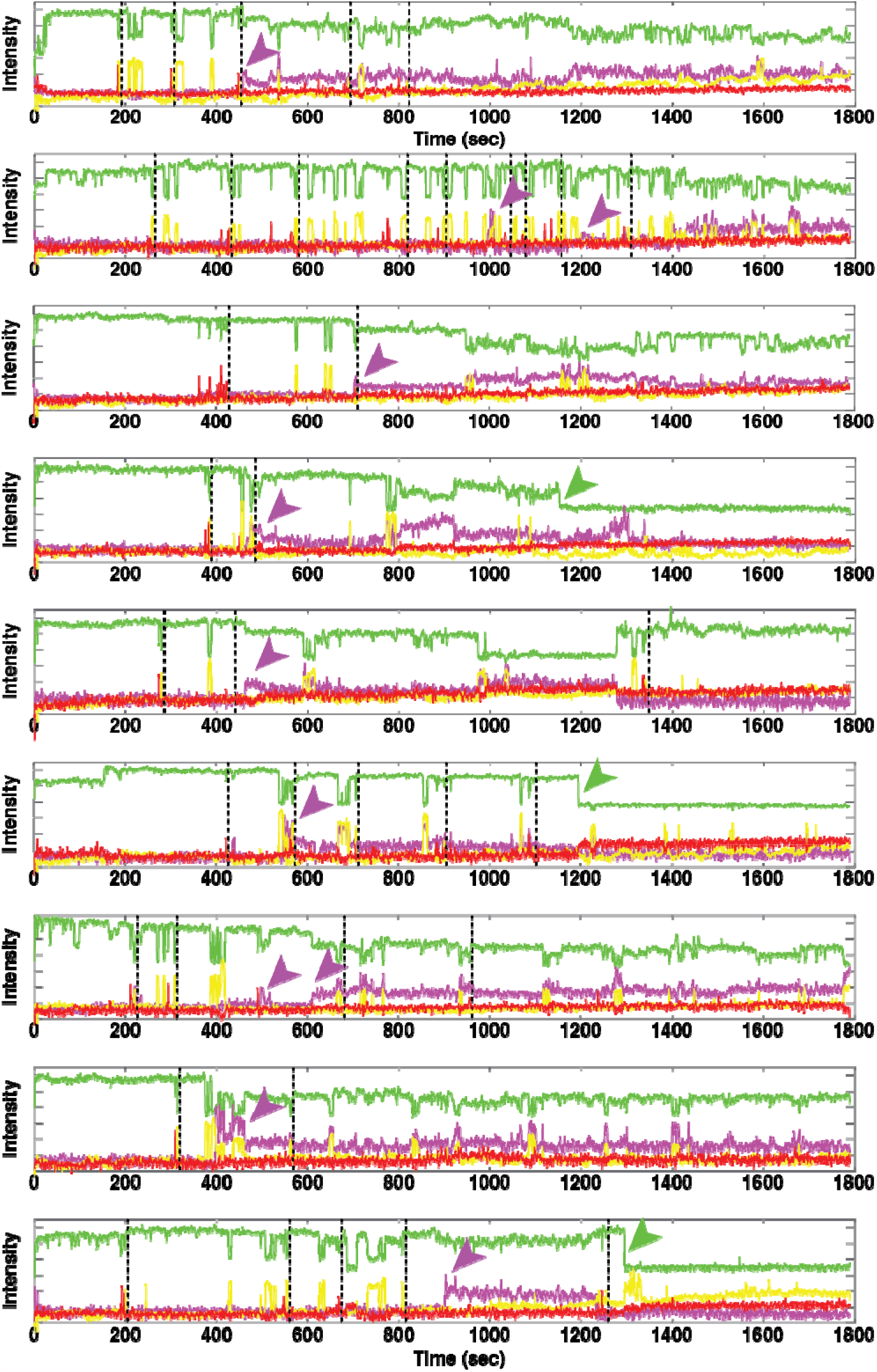
Example traces visualizing real-time telomerase activity by ZMW-FRET in presence of DNA-probe oligonucleotide (magenta). Dashed lines indicate G6 incorporation events preceding a RAP phase, cf. Fig. 3. Loss of LD555 donor fluorescence (green arrows) represents DNA dissociation from telomerase or dye photo-bleaching. An offset is applied to the baseline of LD555 for clarity. The association and onset of DNA-dynamics is indicated by arrowheads in magenta.

**Supplementary Fig. 6.**
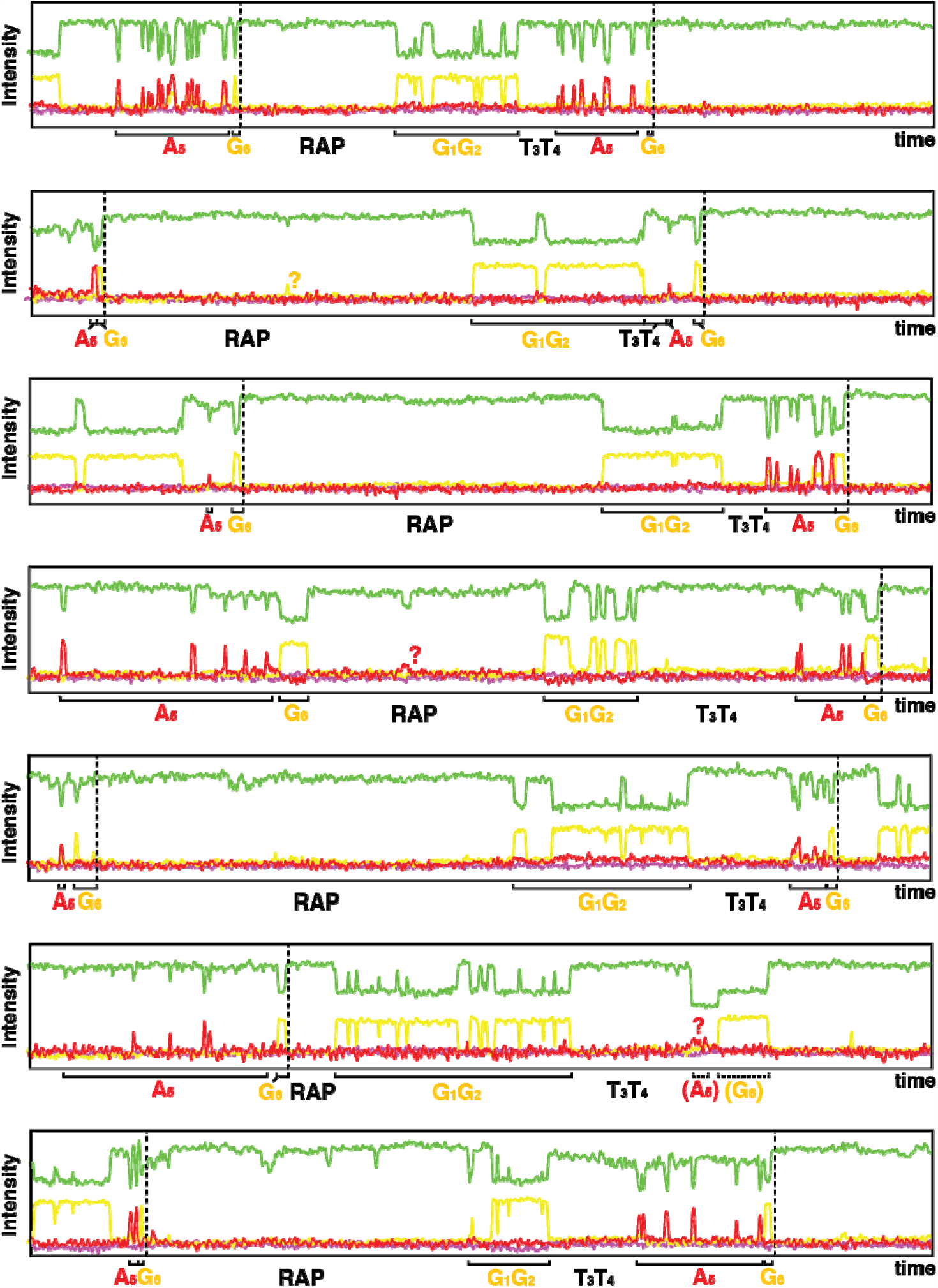
400 second time windows of example traces to highlight the repetitive FRET pattern that corresponds to individual telomeric repeats, cf. Fig. 3b. The probabilistic nature of signal clusters is most prominent for A5 (red) and G1/G2 signals at template positions rU5 and rC1/rC2, respectively. The life-times of G1/G2 clusters exceed expectations for the homo-dimer G1/G2 and may indicate specific RAP steps to be completed after G1-binding, but prior to G1-incorporation, cf. Supplementary Fig. 9.

**Supplementary Fig. 7.**
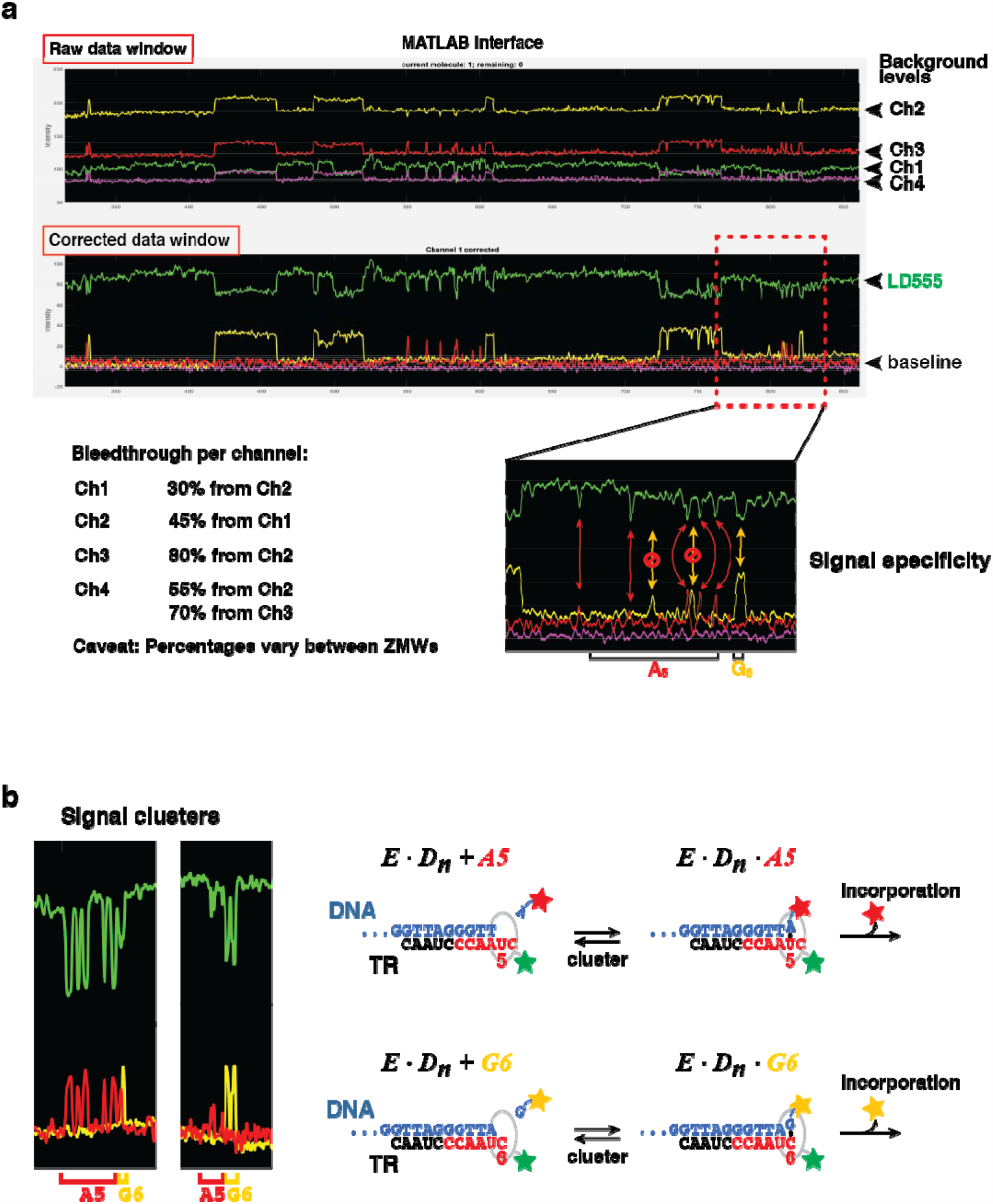
a. Bleed-through correction is performed in Matlab through relative signal subtraction. Percentages of relative bleed-through are given below. Values vary between ZMWs and are adjusted as needed by visual inspection of the respective trace. Enlarged section of the example trace shows anti-correlated (true) FRET events for an A5-cluster followed by a short G6 cluster. Bleed-through correction aids in the visual detection of unspecific Ch2 (yellow) events as highlighted by crossed arrows. **b**. FRET-pulses are heterogeneous and can appear as clusters, indicating nucleotide association and dissociation at a particular template position. Eventual incorporation leads to release of the fluorescence dye after nucleotide hydrolysis and loss of FRET, cf. Fig. 2d.

**Supplementary Fig. 8.**
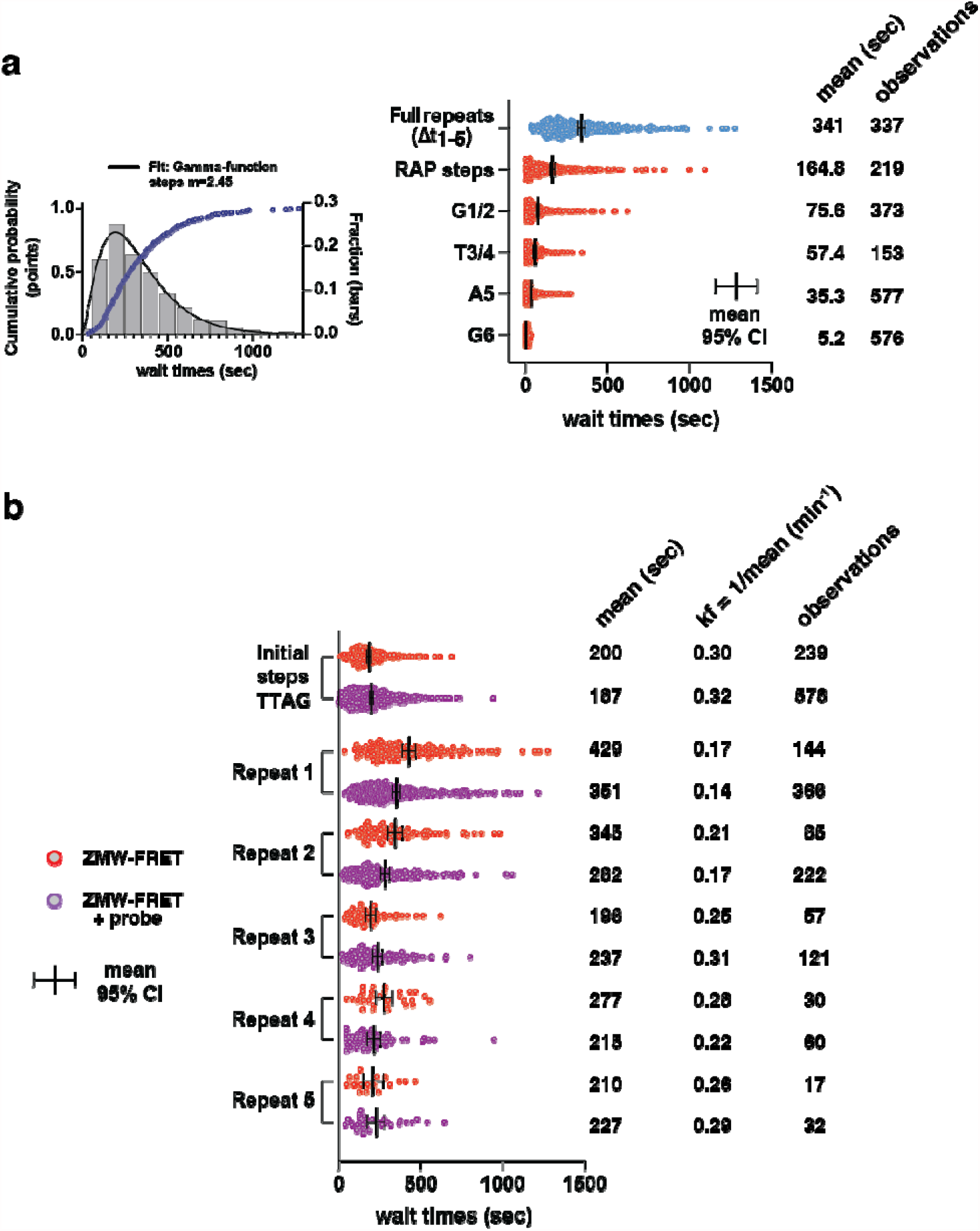
a. Wait time frequency distributions of full repeats and repeat sub-steps, cf. Fig. 3b. The Gamma-function fit (Matlab MEMLET) indicates multiple rate-contributing steps within given repeats. At right, time distributions for the population of full repeats, as well as for repeat sub-steps are given with their mean and 95% confidence interval. **b**. Wait time distributions are given for individual repeats across two independent datasets in absence (red) and presence (magenta) of DNA probe, respectively. Wait times for the initial completion of the telomeric primer (TTAGGG)3 are included. The mean and 95% confidence intervals are indicated. Forward rate constants are calculated as kf = 1/mean to describe an individual repeat as single irreversible step, cf. Fig. 3c.

**Supplementary Fig. 9.**
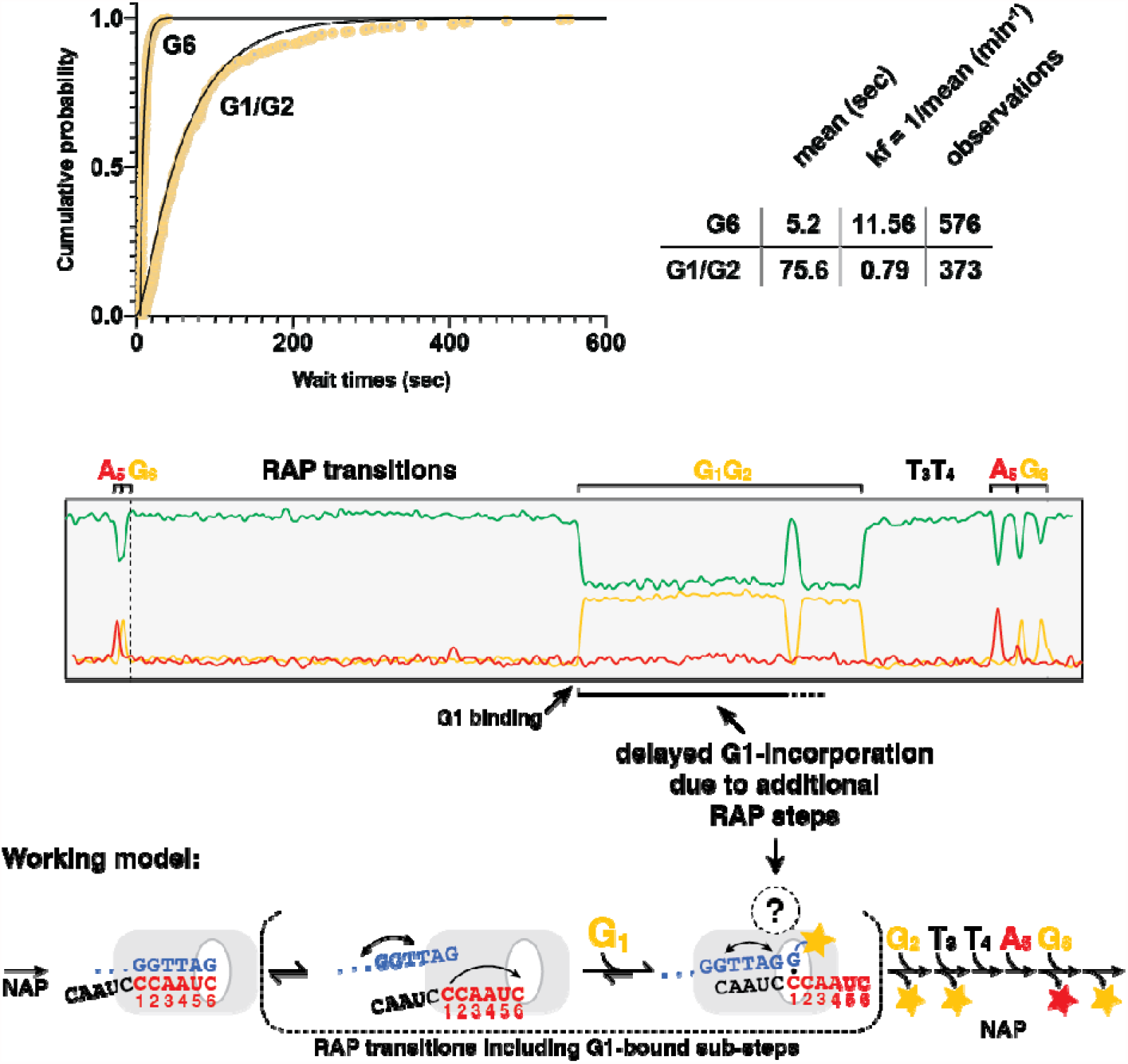
Top panel, comparison of cumulative frequency wait time distributions of G6 and G1/G2 clusters. Single-exponential (G6) and double-exponential (G1/G2) fits are in black. Single-step rate constants are given for conceptual simplification and were calculated as kf = 1/mean. Below, example trace highlighting the prolonged life-times of G1/G2 clusters. The model schematic assigns a potential mechanism of mandatory RAP steps that delay G1-incorporation, as shown in Figure 3b and 3e.

**Supplementary Fig. 10.**
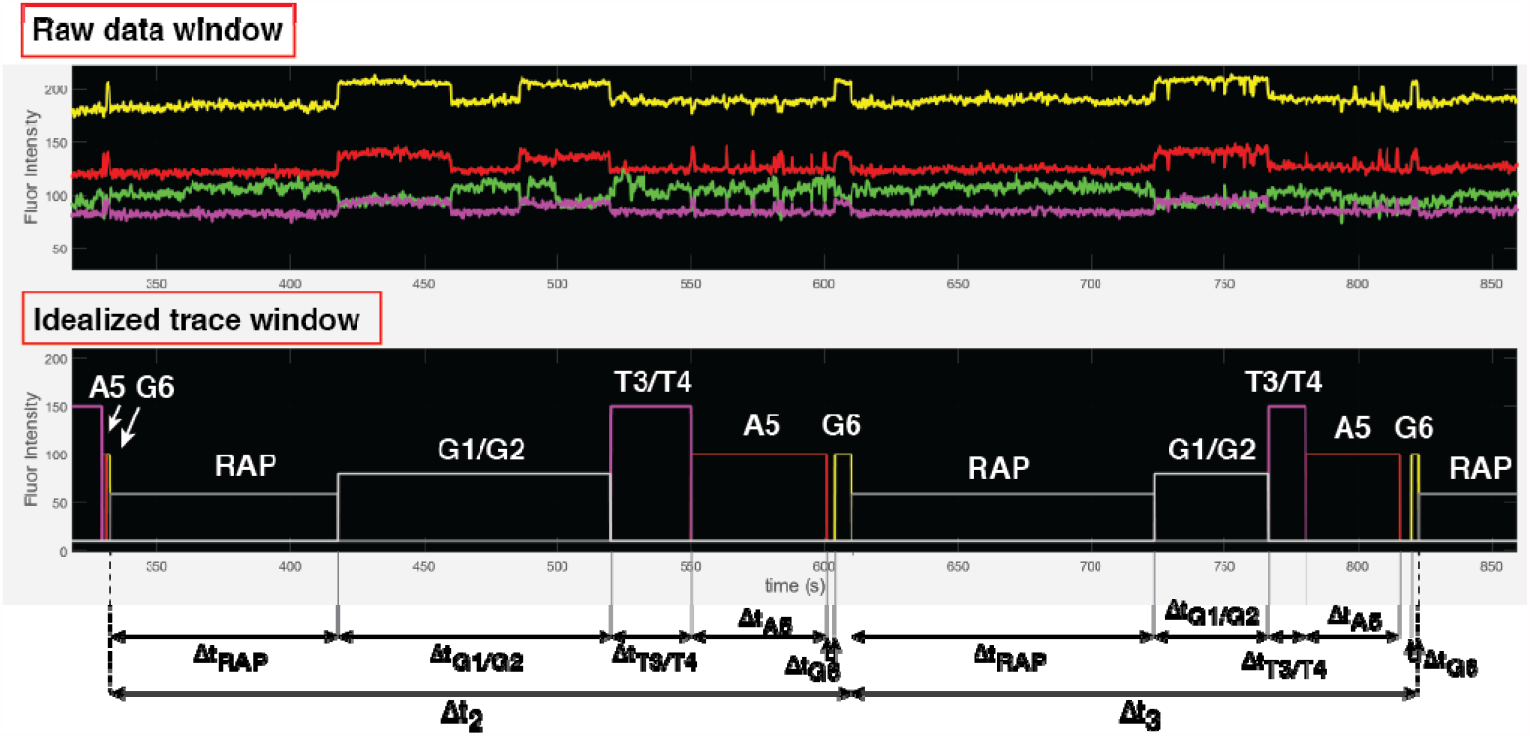
State assignment is performed manually in Matlab and raw data traces are converted to an “idealized state trace”. The amplitudes of states are set to arbitrary values for visual distinction. Individual wait times between state transitions are extracted and analyzed as needed.

**Supplementary Table 1.**
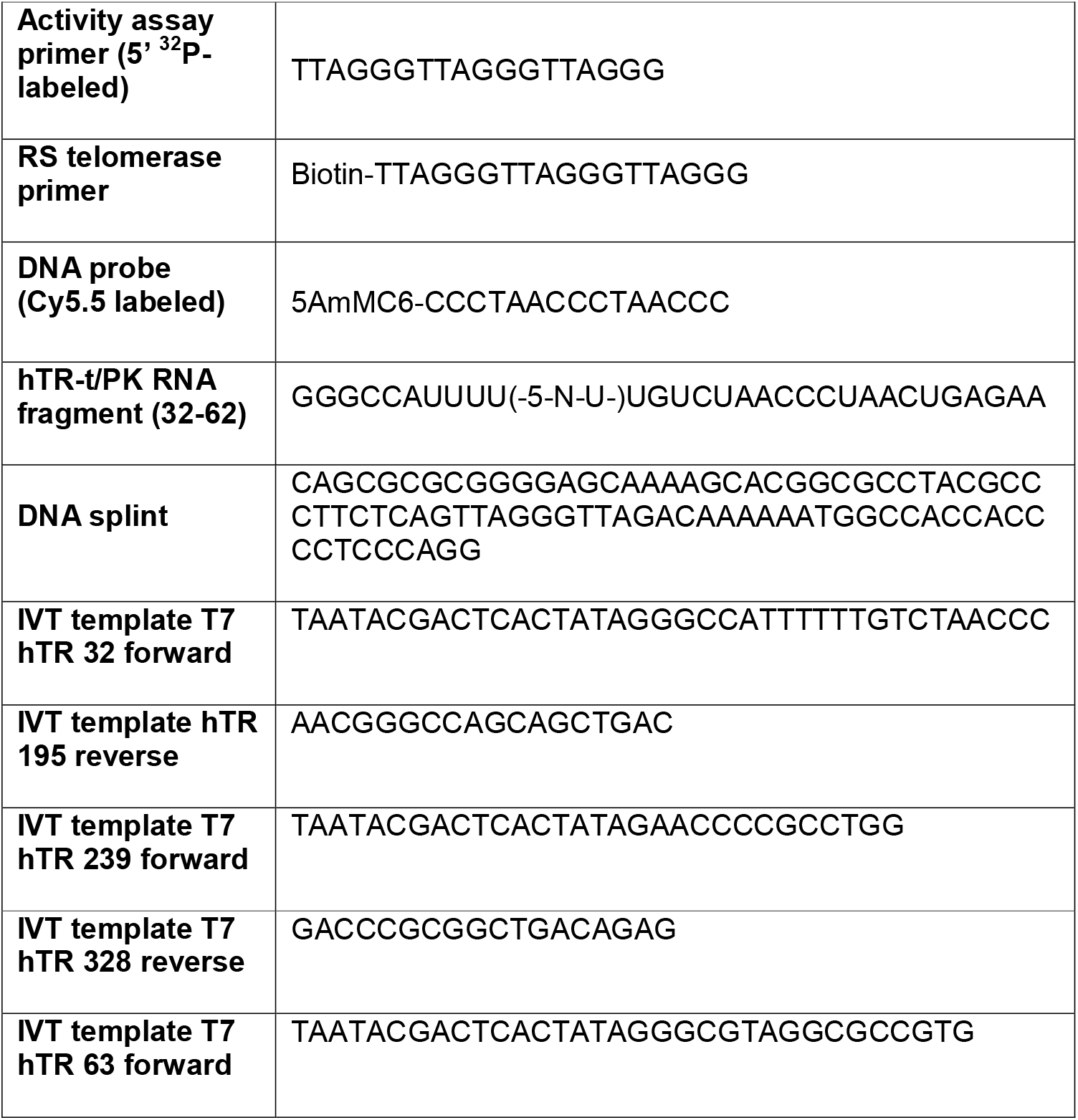
Oligonucleotides used in this study (5′ to 3′). DNA oligonucleotides were purchased from Integrated DNA Technologies, modified RNA fragments from Dharmacon.

## Online Methods

### RNA Preparation

#### RNA in-vitro transcription

In vitro transcription (IVT) DNA templates for hTR CR4/5 (hTR 239-328), hTR template/pseudoknot (hTR t/PK, 32-195 and 63-195) were generated by PCR fusing the T7 promotor to the respective DNA sequence of hTR (Supplementary Table 1). A total of 400 μl PCR reactions were added directly to the IVT mix. IVTs were carried out using homemade T7 RNA polymerase in 5 ml reactions of 40 mM Tris-HCl, pH 8.0, 35 mM MgCl _2_, 0.01% (v/v) Triton X-100, 5 mM DTT, 1 mM spermidine, 2.5 mM each NTP and 40 U RNasin Plus (Promega). The reactions were incubated overnight at 37°C followed by the addition of 10 U TURBO DNase (Thermo Fisher Scientific) and incubation for 15 min at 37°C. The RNA was enriched by isopropanol precipitation and purified by denaturing urea polyacrylamide gel electrophoresis (PAGE). The IVT mix of hTR t/PK 63-195 destined for splinted ligation reactions (see below) was modified to contain 1 mM of each NTP and 5 mM GMP to obtain 5′ monophosphate groups.

#### RNA dye-labeling

To generate dye-labeled telomerase, the synthetic hTR template/pseudoknot (t/PK) fragment 32-62 was purchased from Dharmacon with an internal 5-Aminohexylacrylamino-uridine (5-LC-N-U) at position U42 (Supplementary Table 1). The RNA was deprotected using provided deprotection buffer (Dharmacon) and following manufacturer’s instructions. Deprotected RNA was desalted using G-25 MidiTrap gravity columns (GE Healthcare), followed by ethanol precipitation in presence of 300 mM sodium acetate, pH 5.2. The pellet was resuspended in 100 μl 0.1 M sodium bicarbonate and brought to a concentration of 200 nmol/ml. For dye-coupling, 20 nmol RNA were combined with an equal volume of anhydrous DMSO containing 100 nmol LD555 (Lumidyne Technologies). The mix was incubated in the dark for 2 h at 37°C, desalted using gravity columns, and dried using a vacufuge (Eppendorf). RNA pellets were dissolved in 60 μl 0.1 M triethylamine acetate (TEAA), pH 7.5, and dye-labeled RNA was purified by HPLC in a 0.1 M TEAA to 60% acetonitrile gradient on a reversed phase C8 column (Agilent Technologies).

#### RNA ligation

Synthetic and dye-labeled hTR 32-62 was ligated to in vitro transcribed hTR t/PK 63-195 by splinted RNA ligation. A 200 μl reaction contained 800 pmol of LD555-hTR 32-62, 1600 pmol hTR 63-195, and 1600 pmol DNA splint (Supplementary Table 1) in 0.5x T4 DNA ligase buffer (NEB). The mix was incubated for 5 min at 95°C and for further 10 min at 30°C. 200 μl ligation mix (1.5x T4 DNA ligase buffer, 8000 U T4 DNA ligase (NEB), 2 mM ATP and 200 U RNasin Plus) was added to the reaction and incubated overnight at 30°C. 10 U TURBO DNase were added and incubated for 15 min at 37°C. The RNA was then phenol-chloroform extracted, ethanol precipitated, and purified by urea PAGE.

### Telomerase Reconstitution

Human telomerase was reconstituted in rabbit reticulocyte lysate (RRL, Promega TnT Quick Coupled Transcription/Translation kit) following the manufacturers specifications. Specific to telomerase, 200 μl TnT mix were supplemented with 5 μg of the plasmid pNFLAG-hTERT (REF) and 1 uM of in vitro transcribed (see above) and unlabeled hTR fragments (hTR template/pseudoknot and CR4/5). In reconstitutions of dye-labeled telomerase, the LD555 t/PK fragment (see above) was added at 0.1 uM final concentration owing to lower relative yields after RNA ligation and dye labeling. The reaction mix was incubated for 3 h at 30°C and quenched with 5 μl of 0.5 M EDTA, pH 8.0 for 30 min at room temperature. For immuno-purification of telomerase, beads from 50 ul ANTI-FLAG M2-agarose bead slurry (Sigma-Aldrich) were washed three times by suspension in 750 μl wash buffer (50 mM Tris-HCl, pH 8.3, 3 mM MgCl_2_, 2 mM DTT, 100 mM NaCl), centrifugation for 30 seconds at 2350 rcf, and removal of the supernatant. Beads were subsequently blocked twice for 15 min in blocking buffer (50 mM Tris-HCl, pH 8.3, 3 mM MgCl_2_, 2 mM DTT, 500 μg/ml BSA, 50 μg/ml glycogen, 100 μg/ml yeast tRNA) under end-over-end agitation at 4°C. Blocked beads were collected by centrifugation, resuspended in 200 μl blocking buffer and added to the telomerase reconstitution reaction in RRL. This binding step was left to proceed for 2 h at 4°C under end-over-end agitation. The beads were then pelleted and washed three times in wash buffer containing 300 mM NaCl, followed by three further wash steps in wash buffer containing 100 mM NaCl. Elution of telomerase from the beads was performed by adding 60 μl elution buffer (50 mM Tris-HCl, pH 8.3, 3 mM MgCl_2_, 2 mM DTT, 750 μg/ml 3xFLAG peptide, 20% glycerol) to the beads and incubation for 1h at 4°C under orbital agitation. The mix was then transferred to a Nanosep MF 0.45 uM centrifugal filter and eluate was collected by centrifugation at 10,000 rcf for 1 min. 5 μl aliquots (3 ul in case of dye-labeled telomerase) were prepared in Lo-bind tubes (Eppendorf), flash frozen in liquid nitrogen and stored at - 80°C until use.

### ^32^P -end-labeling of DNA primers

50 pmol of DNA primer was labeled with gamma-^32^P ATP and T4 polynucleotide kinase (NEB) in a 50 ul reaction in 1x PNK buffer (NEB). The mix was incubated for 1 h at 37°C followed by heat inactivation of T4 PNK at 65°C for 20 min. The primers were purified on Centrispin columns (Princeton Separations) and brought to a final concentration of 50 nM in nuclease-free water.

### Primer extension assays

Bulk activity assays were performed using 5 μl immuno-purified telomerase brought to 15 μl reaction volume of 50 mM Tris-HCl, pH 8.3, 50 mM KCl, 3 mM MgCl _2_, 2 mM DTT. ^32^P-end-labeled DNA primers were used at 50 nM and nucleotide concentrations were as indicated (ranging from 0.1 μM to 10 μM of each dATP, dTTP and dGTP). Dye-dG6P and dye-dA6P were used at 3.5 μM and 8 μM, respectively, to be consistent with RS assay conditions. dTTP was used at 10 μM when mixed with dye-dG6P and dye-dA6P. Reactions were incubated for 90 min at 30°C and quenched with 200 μl TES (10 mM Tris-HCl, pH 7.5, 1 mM EDTA, 0.1% SDS). DNA products were phenol-chloroform extracted and ethanol precipitated. DNA pellets were resuspended in formamide gel loading buffer (50 mM Tris Base, 50 mM boric acid, 2 mM EDTA, 80% (v/v) formamide, 0.05% (w/v) each bromophenol blue and xylene cyanol) and resolved on a 12% denaturing urea PAGE gel. The gel was then dried and imaged using a storage phosphor screen and Typhoon scanner (GE Healthcare).

### RS assay

#### Chip preparation and enzyme immobilization

RS chips (SMRT cell, PacBio) were washed three times each with 40 μl TP50 buffer (50 mM Tris-HCl, pH 8.0, 50 mM KCl). 20 μl Neutravidin was supplied to the chip and incubated for 5 min at room temperature. Neutravidin was removed, followed by three wash steps each with 40 μl telomerase buffer (50 mM Tris-HCl, pH 8.3, 50 mM KCl, 3 mM MgCl_2_). 5 nM stocks of biotin-(TTAGGG)3-primer in TLi (50 mM Tris-HCl, pH 8.3, 50 mM LiCl) were heated to 95°C for 3 min to melt primer aggregates or G-rich structures. Primers were cooled on ice. 3 μl immuno-purified LD555-telomerase was combined with 1 μl of 5 nM heat-treated primer and incubated for 20 min at 30°C. 16 ul of telomerase buffer was added to this mix at room temperature. The resulting 20 μl enzyme mix was supplied to an RS chip, sealed with parafilm, and incubated for 15 min at 30°C. The enzyme mix was removed and the chip washed three times each with telomerase buffer.

#### RS setup and chip imaging

The RSII real time DNA sequencer (RS, PacBio) was initialized from the user interface. The green laser (532 nm) was set to a laser power of 0.24 μW/μm^2^. A calibration measurement was previously conducted at the same laser power as a reference according to the manufacturer’s instructions. The movie length was 30 minutes at a frame rate of 10 Hz. The chip clamp temperature (∼ZMW temperature) was 30°C. A dispense protocol was selected for fluidic delivery of 20 μl delivery solution (see below) to the chip at 140 seconds into the movie. After initialization, enzyme-treated chips were supplied with telomerase imaging buffer (50 mM Tris-HCl, pH 8.3, 50 mM KCl, 3 mM MgCl_2_, 2.5% (v/v) N-Formylmorpholine, 3.5 μM dye-dG6P, 8 μM dye-dATP, oxygen-scavenging system [2.5 mM PCA (protocatechuic acid), 2.5 mM TSY, and 2x PCD (protocatechuate-3,4-dioxygenase), purchased from Pacific Bioscience]) and the silica surface of the chip was cleaned with methanol. Chips were then mounted in the SMRT-cell chamber of the RS. Telomerase imaging “delivery” solution (imaging buffer including 20 μM dTTP, and 40 nM DNA probe where indicated) was provided in a 96well sample plate in the sample chamber of the RS and kept at 4°C until delivery. Data acquisition was started from the user interface and four-channel intensity information was written to an .h5 file.^21^

#### Data analysis

Fluorescence trace extraction and data analysis were performed with custom-written Matlab scripts (available upon request). Procedures adhered to the principle work flow described below. ∼150,000 four-color traces were extracted from the RS .h5 movie file sorted by increasing distance from the chip center (script RS_DISTANCE_FILTER). Traces were screened for fluorescence intensity anti-correlation (i.e. FRET) between detection channels 1 (green) and 3 (red), and saved as FRET-positive trace subset (script RS_FRET_picking_auto). The trace subset was manually screened for true FRET events (as opposed to noise or artifacts) using the trace viewer and selector script RS_PICKTRACES. A selected subset was saved as true FRET-positive and used for manual assignment of FRET states using RS_ASSIGNSTATES (Supplementary Fig. 10a). For individual ZMWs, this script plots fluorescence intensity of four channels over a common time axis. Using the ginput() Matlab function, a state of interest was defined by manually selecting two time points corresponding to the beginning and end of the state. Using a custom range of colors and pre-defined state amplitudes, idealized state traces are generated, viewed in real time, and saved in an array (Idealized trace array, ITA). One individual molecule (i.e. ZMW) was viewed and assigned at a time. The HMM array was then used for the extraction of wait times between individual states as desired for single-molecule and kinetic analysis. For that, the diff function was applied to the ITA array and state transitions (and their time values) were identified as non-zero values. Scripts (RS_REPEAT_TIMES, RS_A5_G6, RS_RAP_G1G2, RS_T3T4) were adjusted as needed and state transition wait times were saved in value tables. Wait time distributions were generated and fitted in Matlab MEMLET^23^ or Graphpad Prism. Rate constants for single irreversible kinetic steps were calculated from 1/mean of the respective population of wait times. The 95% CI for wait time distributions was determined in Prism. For visualization and interpretation purposes, raw data traces were corrected for background fluorescence and spectral bleed-through using the script RS_CORRECTION (Supplementary Fig. 7a). For that, fixed relative percentages of intensity were subtracted from each color trace, according to the scheme Ch1corr=Ch1-0.3*Ch2, Ch2corr=Ch2-0.45*Ch1, Ch3corr=Ch3-0.8*Ch2, Ch4corr=Ch4-0.55*Ch2-0.7*Ch3. These values were adjusted as needed after visual inspection of corrected traces.

